# Genetic polymorphism of CYP2C19 and subcortical variability in the human adult brain

**DOI:** 10.1101/2020.12.18.423183

**Authors:** Julia C. Stingl, Catharina Scholl, Julia E. Bosch, Roberto Viviani

## Abstract

Pharmacogenetic studies have shown involvement of cytochrome P450 enzymes in the metabolism of psychotropic drugs. However, expression and activity on endogenous substrates in the brain may underlie a constitutive role of these enzymes beyond drug metabolism. CYP2C19, which is expressed in the human fetal brain during neurodevelopment, shows affinity for endogenous compounds including monoaminergic neurotransmitters, steroid hormones, and endocannabinoids. In this study (N=608), we looked at the genetic polymorphism of CYP2C19 and its potential associations with structural phenotypes of subcortical brain volume with structural imaging. Using two independent volume estimation techniques, we found converging evidence for a positive association between *CYP2C19* activity scores, as inferred from the genotype, and basal ganglia and hippocampal volume. This association was present only in female individuals, raising the possibility that effects on brain morphology may arise through a mechanism involving the metabolism of estrogen steroids.

## Introduction

CYP2C19 is a genetically polymorphic drug-metabolizing enzyme that, due to broad substrate specificity, metabolizes different classes of drugs including several psychotropic drugs such as antidepressants (selective serotonin-reuptake inhibitors, tricyclic antidepressants), benzodiazepines, and antiepileptics. The genetic polymorphism influences the range in CYP2C19 enzyme activity in individuals from complete loss of functionality (poor metabolizer phenotype, *CYP2C19*2/*2* carriers) to severalfold enhancement (ultrarapid metabolizers, *CYP2C19*17* carriers, Sim et al. 2006). Beside its role in xenobiotic metabolism, CYP2C19 is also involved in biotransformation of endogenous compounds such as neurotransmitters (e.g., hydroxytryptamine), steroid hormones (estrogen, progesterone, testosterone), fatty acids (cannabidiol) and arachidonic acid (Stingl et al. 2013; Persson et al. 2013). While not highly expressed in extrahepatic tissues of the adult, transient CYP2C19 expression in the human fetal brain may play a role during embryogenesis and neurogenesis (Ingelman-Sundberg et al. 2014).

An important approach to documenting the role of CYP2C19 on brain structure and function is to look at individual differences associated with its genetic polymorphism. In a study of the functional phenotype of the *CYP2C19*17* genotype predicting ultrarapid metabolic activity, elevated CYP2C19 expression was associated with depressive symptoms and higher tendencies to suicidal impulses in humans (Jukic et al. 2016). Furthermore, the poor metabolizer phenotype was shown in the same study to be associated with reduced volume of the hippocampus, a common finding in studies of depression (Sheline 2011). In transgenic mice expressing the human *CYP2C19* gene, impairment of hippocampal serotonin and BDNF homeostasis and increased anxiety has been shown to be involved in an anxious depressive phenotype (Persson et al. 2013; Jukic et al. 2016).

The mechanism behind the possible effects of *CYP2C19* genotype on the brain, however, are not fully understood. An intriguing aspect of this issue is that CYP2C19 is not expressed or active as enzyme in the adult brain and in neuronal cells. However, CYP2C19 mRNA was detected during neurodevelopment in neuronal cells, a finding that has led to several hypotheses on the possible endogenous substrates of CYP2C19 involved in neurogenesis (Ingelman-Sundberg et al. 2014). CYP2C19-mediated metabolism of cannabinoid compunds to the 6-alpha- and the 7-hydroxy-metabolites (Jiang et al. 2011) may affect cannabinoid signaling in the developing brain (Ingelman-Sundberg et al. 2014; Berghuis et al. 2007), as in the migration of GABA-ergic neurons to the hippocampus (Berghuis et al. 2005). Also the affinity of CYP2C19 for steroid hormones (including endogenous estrogens such as estradiol (Cheng et al. 2001; Cribb et al. 2006) deserves to be brought to attention because of the large quantitative effects of sex on brain structure and the prominent role of estrogens in controlling sexual dimorphism in the fetal brain, where estradiol is also synthetized and metabolized locally (McCarthy 2008).

In the present study, we looked at the genetic polymorphism of CYP2C19 and its potential associations with structural subcortical brain imaging phenotypes. While compatible with the findings of Jukic et al. (2016), our data showed a more extensive involvement of subcortical volumes, which increased in association with higher *CYP2C19* activity scores. Our data were also suggestive of an impact of *CYP2C19* genotype on sexual dimorphism, pointing to a possible mechanism through which these effects are produced during brain development.

Methodologically, we verified the robustness of volumetric estimates of the hippocampi and other subcortical structures by employing two very different techniques: the first was the same used by Jukic et al. (2016), based on segmenting subcortical volumes directly; the second was based on the amount of deformation required to register these structures to a common template (Ashburner et al. 1998; Gaser et al. 1999). These two different approaches provided convergent assessments of the effect of *CYP2C19* genotype on subcortical volumes.

## Methods

The study protocol was approved by the local ethics committees in Ulm and Bonn and the study was registered by the German Clinical Trials Register (DRKS-ID: 00011722, BrainCYP Study).

### Participants

The sample used in the study consisted of N = 608 healthy individuals obtained by pooling data from three samples (Allegra Ulm, N = 175, BrainCYP Ulm, N = 378, BrainCYP Bonn, N = 55). The final sample included 342 females (average age 24.1, std. deviation 4.3).

Healthy participants were recruited primarily from the local universities and by local announcements. Exclusion criteria were neurological or medical conditions, use of medication (except hormonal contraceptives and L-thyroxin), alcohol or drug abuse or a history of mental illness. In order to minimize the risk of ethnic stratification, Caucasian descent was ascertained by asking about the genetic background of both parents. For more detail on cohort characteristics see Table 1.

**Table 1.** Genetic composition of the sample.

| Genotype | Activity score | N | N females | N males |
| --- | --- | --- | --- | --- |
| *2/*2 (PM) | 0 | 10 | 7 | 3 |
| *2/*1 | 1 | 113 | 59 | 54 |
| *2/*17 | 1 | 22 | 15 | 7 |
| *1/*1 (WT) | 2 | 263 | 156 | 107 |
| *1/*17 | 3 | 167 | 89 | 78 |
| *17/*17 (UM) | 4 | 33 | 16 | 17 |

PM: poor metabolizer; WT: wild type; UM: ultrarapid metabolizer

### Genotyping

As a part of the study procedures, a 7ml EDTA-blood sample was taken from all participants. For genotyping, genomic DNA was isolated from the biosamples using magnetic beads with the MagNA PureLC DNA Isolation Kit—Large Volume (Roche Diagnostics GmbH., Mannheim, Germany). Genotypes for *CYP2C19*, rs12248560 (C>T) (*CYP2C19*17*) and rs42442085 (G>A) (*CYP2C19*2*), were determined by real-time PCR using SimpleProbe® probes (LightSNiP assays), designed by TIB Molbiol (Berlin, Germany). To this end, 50 ng DNA were amplified using Lightcycler® FastStart DNA Master HybProbe Mix (Roche, Germany) and the respective LightSNiP Assay. The following PCR protocol was applied: 10 min at 95°C, followed by 45 cycles for amplification: 10 s at 95°C, 10 s at 60°C, 15 s at 72°C. Melting curve analysis was performed to distinguish between the different genotypes. We used a goodness-of-fit c2 test to evaluate agreement with Hardy–Weinberg expectations. *CYP2C19* diplotypes were assigned activity scores as shown in Table 1. If no mutation for rs12248560 and rs42442085was detected, wildtype carrier status was assumed.

### MRI data collection and analysis

The first sample was collected with a Siemens Allegra 3-Tesla scanner on the premises of the Department of Psychiatry of the University of Ulm, Germany, as previously described in Viviani et al. (2007). Briefly, MPRAGE images were collected with size 256×256×208 voxels, voxel size 1×1×1 mm, TR/TE 3300/96, flip angle 90°, bandwidth 1220 Hz/pixel, and field of view of 240×240. The second sample was collected on the same premises from a 3-Tesla Siemens Prisma scanner, while the third sample was collected at the DZNE in Bonn, Germany with a 3-Tesla Skyra scanner using the same sequence on both sites (as previously described in Viviani et al. 2017). In brief, multi-echo MPRAGE images were collected with size 256×256×176, voxel size 1×1×1 mm, TR/TE 2500/1.48, 2.98, 4.48, 5.98, 7.48, flip angle 7°, bandwidth 780 Hz/pixel, and field of view of 256×256.

FIRST is a software package for the estimation the volumes of a set of subcortical structures (hippocampus, amygdala, caudate, nucleus accumbens, pallidum, putamen, thalamus, and brainstem) from T1-weighted structural images using a Bayesian approach (Patenaude et al. 2011). The FIRST code is part of the FSL software package for the analysis of imaging data (https://fsl.fmrib.ox.ac.uk/fsl/fslwiki/FIRST). We used the default settings in this package to estimate volumes of these structures. FIRST computes a segmentation of these subcortical structures and counts the number of voxels in each of the segments. In the Results section, we refer to the count for all these structures as “subcortical volume”, a proxy for a global volumetric effect in FIRST analyses.

Deformation-based morphometry analyses were conducted with the SPM12 package (https://www.fil.ion.ucl.ac.uk/spm/) using the DARTEL diffeomorphic registration algorithm (Ashburner 2007) after conducting a preliminary segmentation with the default settings provided in this package. This preliminary segmentation was also used to derive cranial and brain volume measures. The DARTEL procedure was subsequently applied to maps of grey and white matter intensities from the segmentation. To obtain estimates of subcortical volumes comparable to those obtained with FIRST, we preliminarily generated masks of subcortical structures by applying FIRST to segment the mean DARTEL-registered image. Since this image is the average from the whole sample, it includes no information on volumetric individual differences. Geometric means of the Jacobians of the deformation computed with DARTEL in each voxel were then computed for each of the subcortical masks. The geometric mean of the Jacobians for the voxels in all subcortical masks gave an estimate of “subcortical volume”.

Statistical analysis of volumetric data was conducted with the freely available statistical software R (version 3.5.2, The R Project for Statistical Computing, https://www.r-project.org/) on the logarithm of volume estimates (analyses on the data without taking the logarithms gave comparable results). Boxplots were drawn with the package ggplot2 by Hadley Wickham (https://github.com/tidyverse/ggplot2).

Voxel-based analyses were conducted using a permutation technique to estimate significance levels corrected for the region of interest of the hippocampus (8000 resamples, Holmes et al. 1996) after smoothing the Jacobian images outputted by DARTEL with an isotropic smoothing kernel FWHM 6mm with the SPM12 software. Clusters were defined by the *p* < 0.01 threshold, uncorrected. In the text, *k* denotes the number of voxels composing the cluster. Overlays were drawn with the freely available display software MRIcron by Chris Rorden (http://www.mccauslandcenter.sc.edu/crnl/mricro).

## Results

Jukic et al. (2016) had shown in two independent samples that CYP2C19 poor metabolizers (homozygous carriers of the *2 allele) had a larger hippocampus than carriers of other genotypes after adjusting for subcortical volumetric effects and the confounds of sex, age and scan hardware versions (for the sake of comparability, the term ‘subcortical volumes’ here refers to the volumes for which the package used by Jukic et al. 2016 provides a volumetric estimate, as detailed in the Methods). In our data and using the same methodological approach of Jukic et al. (2016) to assess volumes, the hippocampal volume of poor CYP2C19 metabolizers was 2.5% larger than in the rest of the sample after adjusting for total subcortical volume, age, sex, and scanning site, but this effect failed to reach significance (*t*_601_ = 1.04, *p* = 0.15, one-tailed, Figure 1). As detailed in the Methods, to verify robustness of volumetric estimates we conducted parallel analyses using a deformation-based method. In the deformation-based analysis of the same model, the effect of poor metabolizer status on hippocampal volume was significant (*t*_601_ = 1.71, *p* = 0.044, one-tailed). However, we noticed that the poor metabolizer phenotype was also weakly associated with subcortical volume as a whole (*t*_602_ = -1.46, *p* = 0.14), one of the adjustment terms in the model.

**Figure 1.**
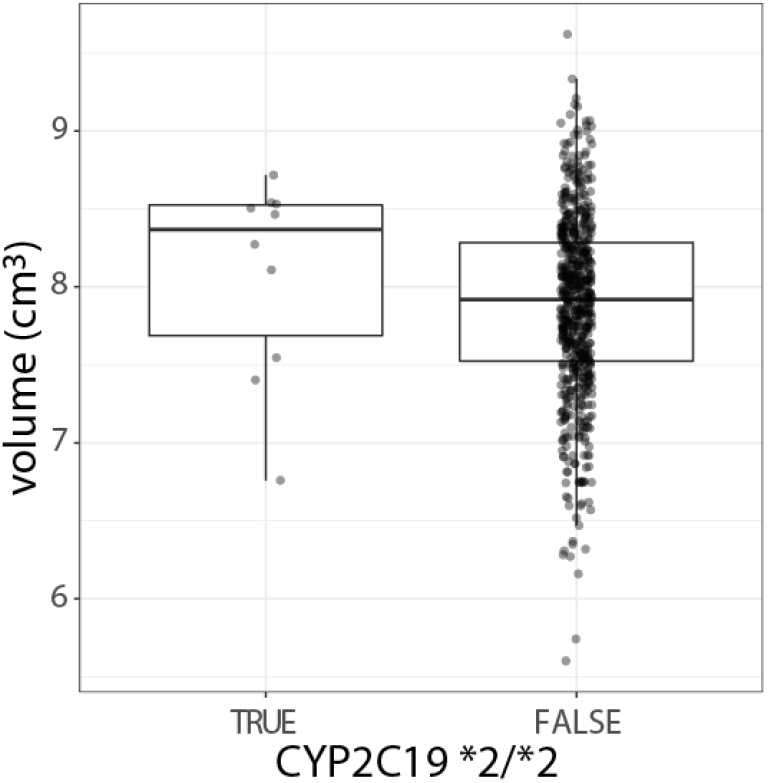
Boxplots of estimated hippocampal volumes, after adjustment for subcortical volume, sex, age and site. The position of the observations on the x-axis was jittered to improve visualization.

When considering subcortical volume in its own right, our data gave a different picture of the relationship between *CYP2C19* genotype and brain structure. This association was significant when tested on the activity score as predicted by the genotype (i.e., when accounting for the effect of all genotypes: *t*_602_ = 2.24, *p* = 0.025; similar results were obtained with deformation-based technique, *t*_602_ = 1.82, *p* = 0.07). A unit difference in the activity score corresponded to an increment of about 0.8% in subcortical volume (giving a difference of 3.2% for the score range; Figure 2).

**Figure 2.**
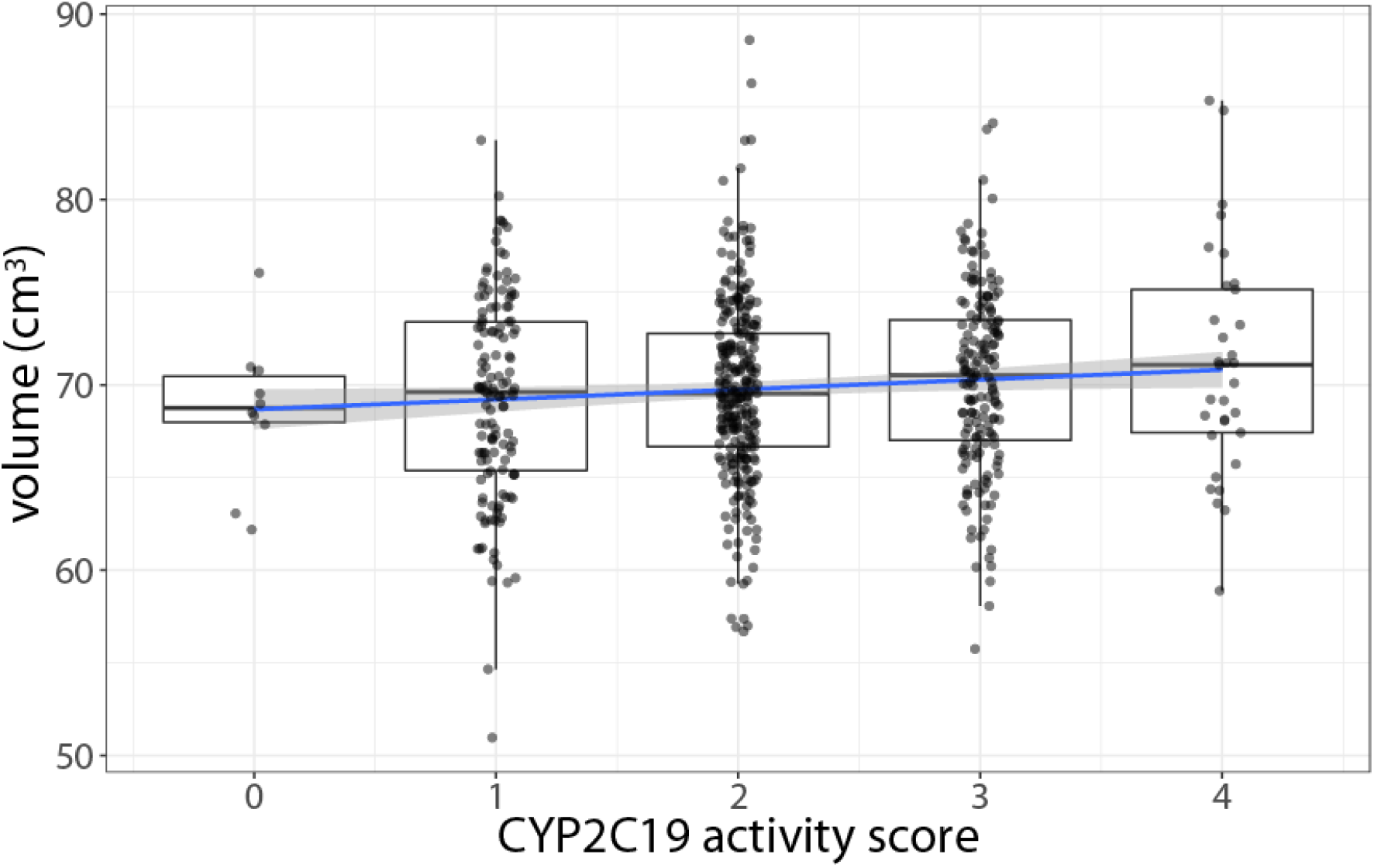
Global subcortical volumes with respect of *CYP2C19* activity scores, adjusted for sex, age, and site. In contrast to Figure 1, voumes increase with activity scores.

This is of potential relevance because, together with the strong association between total subcortical volume and hippocampal volume (more than 90% in our data), it may affect estimates of the association between this latter and the ultrarapid metabolizer phenotype when adjusting for subcortical volume. Indeed, in a model without adjustment for subcortical volume, hippocampal volume was in our data positively associated with the *CYP2C19* activity score (1% volume increment per activity score unit, *t*_602_ = 2.28, *p* = 0.023).

As one might surmise by these findings, in the absence of adjustment for subcortical volume the positive association between *CYP2C19* activity scores and volume was present in most subcortical structures (see Table 2). Because the precision with which the volumes of anatomical structures are estimated varies, significance values do not necessarily correspond to the estimated size of effects.

**Table 2.**
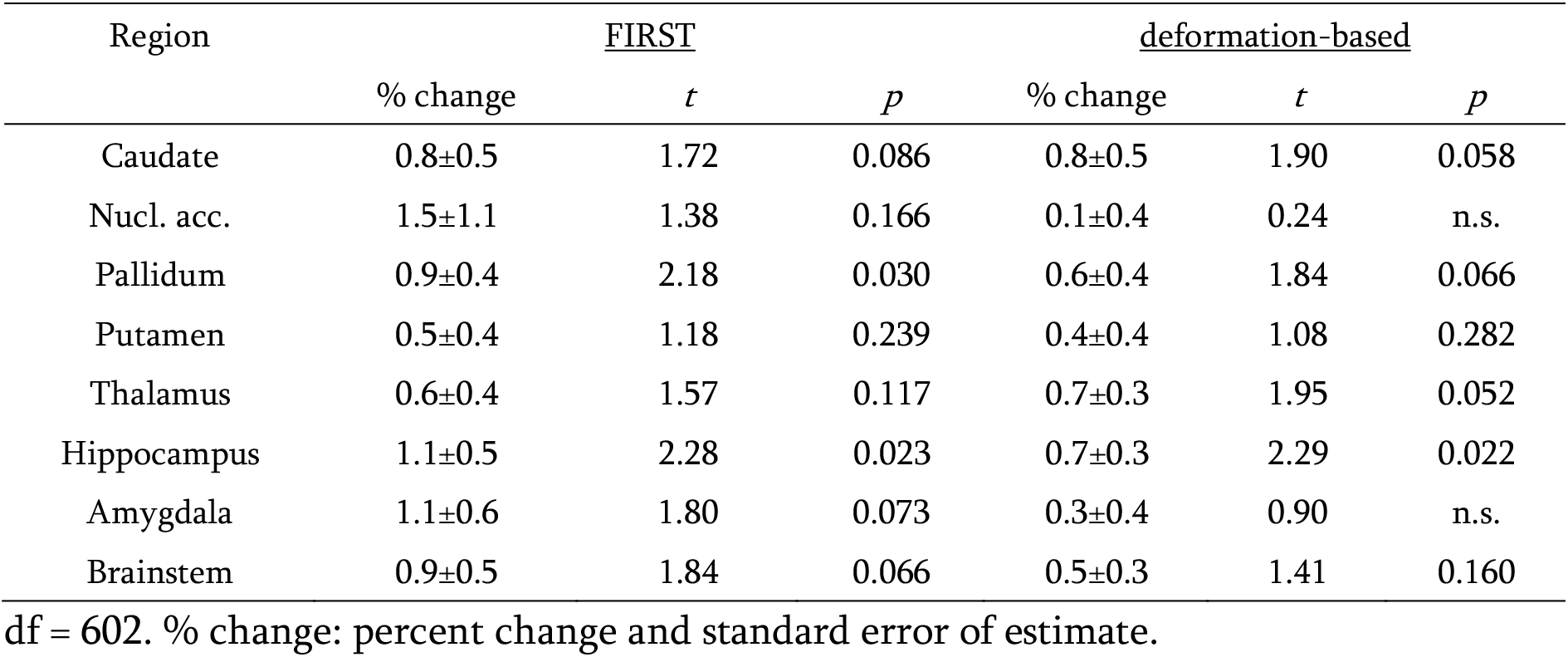
Effect of *CYP2C19* activity score on subcortical volumes

The effects of CYP2C19 polymorphism on cranial or brain volume were less marked than those for subcortical volumes (cranial volume, 0.5% increase per score unit, *t*_602_ = 1.57, *p* = 0.12; brain volume, 0.3%, *t*_602_ = 1.06, n.s.), suggesting some degree of specificity of the effect on subcortical volumes. After adjusting for subcortical volume, there was no residual effect on the effect on cranial and brain volumes, which again suggests that the small effects on cranial and brain volume may be driven by the increase in the subcortical compartment. In line with this interpretation of the data, the effects *CYP2C19* activity scores on the subcortical volume persisted when adjusting for brain (0.5% per unit score increase, *t*_601_ = 2.23, *p* = 0.02) or cranial volume (0.4%, *t*_601_ = 1.60, *p* = 0.06, one-tailed). In summary, the effects of *CYP2C19* activity scores on subcortical volumes appeared to be specific and decoupled from effects (or lack thereof) on brain or cranial size.

### Sex differences

Because of the involvement of CYP2C19 in the biotransformation of steroid hormones and estrogens, we were interested in looking at whether the effect of the genetic polymorphism was sensitive to sex differences. In our data, the association with subcortical volume was stronger in females (*t*_337_ = 2.74, *p* = 0.003), where one unit score corresponded to about 1.2% relative increment, than in males (0.2% increase per score unit, *t*_261_ = 0.41, n.s.), as shown in the interaction (*t*_601_ = 1.51, *p* = 0.065, one-tailed).

As in the data for the whole cohort, this interaction could not be attributed to a generic effect on cranial size reflected in the data for the subcortical volume. After adjusting for cranial volume, the interaction between subcortical volume, sex, and *CYP2C19* activity score persisted (*t*_600_ = 1.69, *p* = 0.046; Figure 3). The deformation-based analysis gave very similar results (*t*_600_ = 1.68, *p* = 0.046).

**Figure 3.**
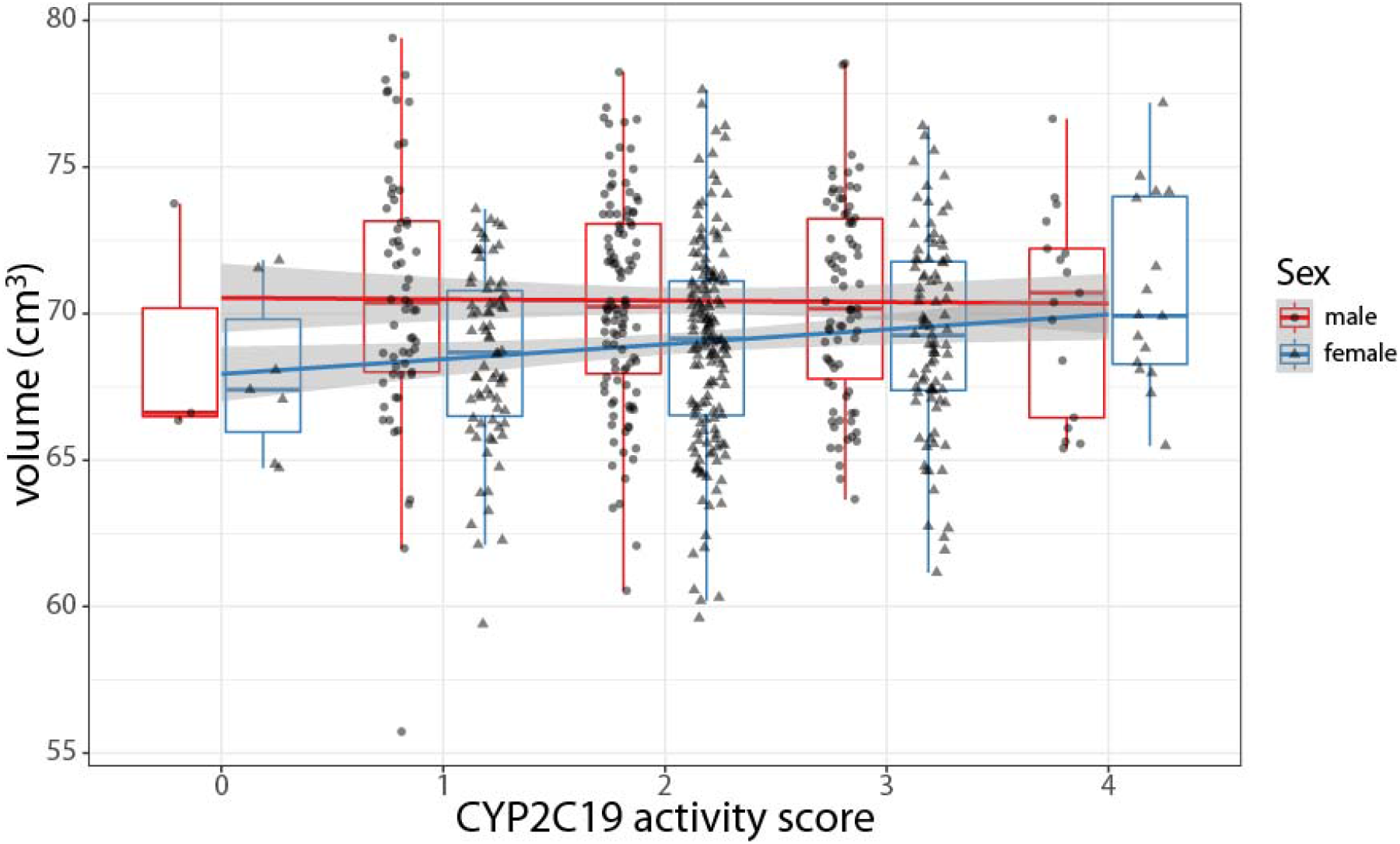
Effects of *CYP2C19* activity on subcortical brain volume in males and females. The regression line for males runs essentially flat (in red), whereas the line for females shows an increase in volume with increasing activity scores (in light blue), so that sexual dimorphism is small or nonexistent in ultrarapid metabolizers (score = 4). Data adjusted for age, site and cranial volume.

Visualization of the sex differences using the deformation-based data revealed a diffuse subcortical involvement in females, with more marked effects in the anterior pallidum and the thalamus (Figure 4).

**Figure 4.**
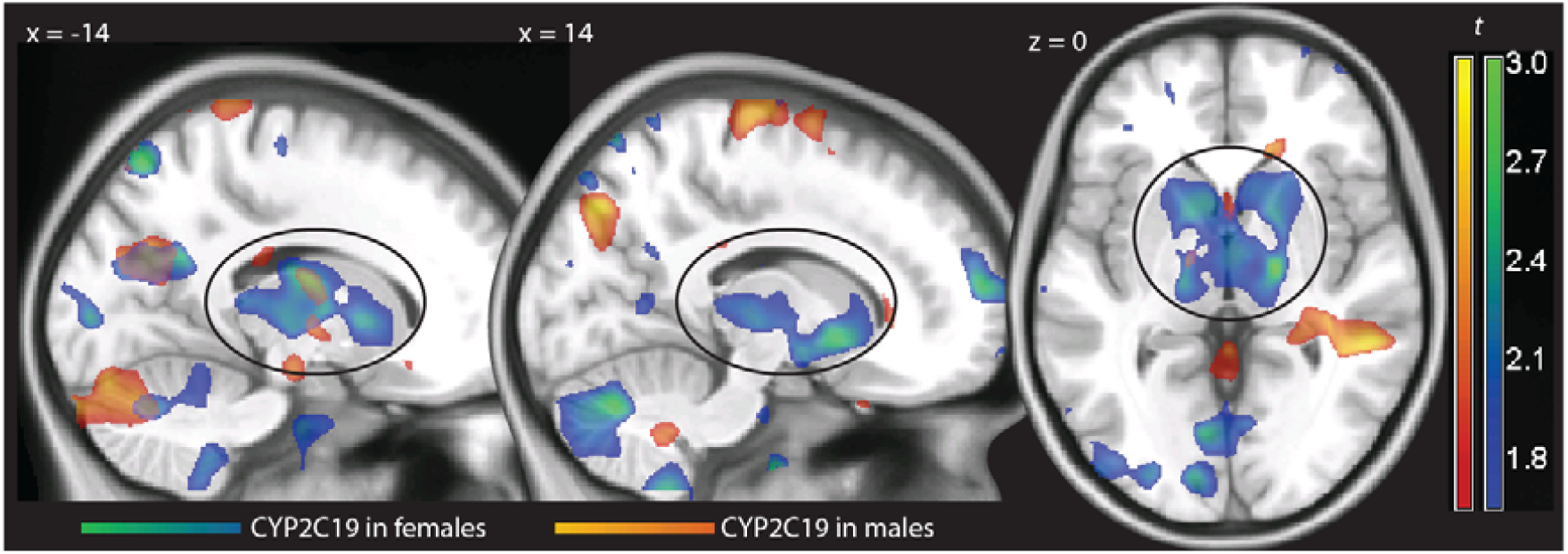
Statistical maps of *t* values obtained from the voxelwise regression of deformation values on *CYP2C19* activity scores in females (in blue-green) and males (red-yellow), thresholded for display purposes at *p* < 0.05, uncorrected, overlaid on a template brain. Positive *t* values indicate larger structures. The ovals outline the effects on the basal ganglia.

## Discussion

In this study of N = 608 healthy individuals we detected a significant association between CYP2C19 activity predicted by genotype and subcortical brain volume. This finding is consistent with previous reports on the existence of effects of *CYP2C19* polymorphism and structural neuroimaging data (Jukic et al. 2016; Stingl et al. 2016). However, our analysis disclosed a more complex set of effects than previously surmised, for which three possible interpretations may be given.

First, while of only borderline significance, our data essentially replicated the finding reported by Jukic et al. (2016) of larger hippocampal size in poor CYP2C19 metabolizers after adjusting for subcortical volume, and showed this effect to be robust relative to the specific technique used to assess hippocampal volume. This finding has been interpreted in support for an affective vulnerability phenotype of CYP2C19 poor metabolizers (Persson et al. 2013). In our data, however, subcortical volume was itself associated with *CYP2C19* activity scores; without adjustment, the association was in the other direction. This suggests that adjustment for subcortical volume may affect the relationship between individual structures and CYP2C19 polymorphism, rather than simply adjusting for a neutral baseline, thus accounting for the reversal of the sign of the association with and without adjustment.

A second interpretive line arises from noting that in our sample the CYP2C19 polymorphism effect on subcortical volume was more pronounced in women. The evidence for the interaction with sex was only suggestive in our data. However, it provides a potential mechanism for the effect of this polymorphism on brain size, because sex alone has a large effect on brain size (Gur et al. 1999). This effect is thought to be mediated by the neurotrophic activity of sex hormones during gestation and adolescence (Goldstein et al. 2001; Lenroot and Giedd 2010). A distinction between sexual dimorphism in brain structure is usually made between a global effect, with ∼10% larger brain size in males, and relative effects that affect most markedly the medial temporal lobe and the basal ganglia (Rijpkema et al. 2012; Lenroot and Giedd 2010). In our sample, the interaction between *CYP2C19* genotype and sex involved subcortical structures, while there was no evidence for a global effect on brain volume.

Several findings support the role of sex hormones in the effects of CYP2C19 polymorphism on brain structure. The role of CYP2C19 in estradiol biotransformation is important for endogenous as well as exogenous estrogens such as oral contraceptives. 17-beta-hydroxydehydrogenation and 16-alpha-hydroxylation of estradiol and similar compound is mediated by CYP2C19 (Cheng et al. 2001; Cribb et al. 2006). Therefore, lower activity of CYP2C19 may cause higher estrogen levels, and vice versa. Consistently with the notion that estrogen exposure is lower in carriers of the high activity *CYP2C19*17* allele, ultrarapid metabolizers have been shown to present lower incidence of breast cancer, and lower recurrences of hormonal sensitive breast cancer after treatment with tamoxifen (Schroth et al. 2007; Justenhoven et al. 2009). Furthermore, the expression of CYP2C19 is partially regulated through a mechanism elucidated recently. The estrogen receptor alpha specifically binds to the promotor of the *CYP2C19* gene and leads to a decrease in enzyme activity through transcriptional down-regulation of *CYP2C19* gene expression, rather than direct estrogen mediated steroid-protein interaction (Mwinyi et al. 2010). Interestingly, the activity of the *CYP2C19*17* allele, which leads to ultrarapid metabolizer phenotypes by an increased transcription rate, is not affected by estrogen receptor alpha-mediated CYP2C19 inhibition as shown in vivo with oral contraceptives (Pedersen et al. 2013). In normal and intermediate metabolizers, estrogen receptor-mediated CYP2C19 inhibition may lead to lower estrogen degradation, shifting activity towards the levels of poor metabolizers. In contrast, in carriers of the *CYP2C19*17* allele, the ultrarapid metabolizer phenotype remains unaffected by estrogen receptor alpha-mediated CYP2C19 inhibition. Thus, the interaction with estrogen-mediated inhibition of the enzyme and estradiol biotransformation as a substrate of CYP2C19 may amplify the phenotypic expression of CYP2C19 genotypes in women. Hence, the association of subcortical volume with *CYP2C19* activity scores in females may derive from overall estrogen exposure that is higher in poor metabolizers leading to an enhanced female dimorphism brain structural phenotype. In contrast, female *CYP2C19*17/*17* carriers were like males with regard to the effects of this genotype on subcortical volume and hippocampus size.

Finally, the observed association between *CYP2C19* genotype and subcortical volume affected specifically the basal ganglia. The function of these subcortical structures is predominantly dopaminergic with effects on reward or overall emotional processing. This hypothesis is consistent with the previously reported higher concentration of dopamine in the brain of transgenic mice expressing human CYP2C19 (Stingl et al. 2016). Several studies have documented sexual dimorphism in humans in basal ganglia dopamine function (Pohjalainen et al. 1998; Laakso et al. 2002; Munro et al. 2009); hence, this finding is not incompatible with the two previous hypotheses.

On a methodological note, we could show that volumetric assessment through MRI led to comparable results across studies irrespective of methodology. The apparent discrepancies in assessing relative effects of individual brain structures were due to issues intrinsic to the rationale for adjusting for global volumetric effects and not to methodology.

## Acknowledgments

This work was supported in part by a Neuron-ERANET grant (project ‘BrainCYP’, grant number BMBF 01EW1402B) and by collaborative grants from the Federal Institute for Drugs and Medical Devices (BfArM, Bonn, Germany, Grants No. V-15981/68502/2014-2017 and V-17568/68502/2017-2020). The authors declare no conflict of interest.

## References

Ashburner, J., 2007. A fast diffeomorphic image registration algorithm. NeuroImage 38, 95–113.

Ashburner, J., Hutton, C., Frackowiak, R., Johnsrude, I., Price, C., Friston, K., 1998. Identifying global anatomic differences: Deformation-based morphometry. Hum. Br. Mapping 6, 348–357.

Berghuis, P., Dobszai, M.B., Wang, X., Spano, S., Ledda, F., Sousa, K.M., Schulte, G., Ernfors, P., Mackie, K., Paratcha, G., Hurd, Y.L., Harkani, T., 2005. Endocannabinoids regulate interneuron migration and morphogenesis by transactivating the TrkB receptor. Proc. Natl Acad. Sci. USA 102, 19115–19120.

Berghuis, P., Rajnicek, A.M., Morozov, Y.M., Ross, R.A., Mulder, J., Urbán, G.M., Monory, K., Marsicano, G., Matteoli, M., Canty, A., Irving, A.J., Katona, I., Yanagawa, Y., Rakic, P., Lutz, B., Mackie, K., Harkani, T., 2007. Hardwiring the brain: Endocannabinoids shape neuronal connectivity. Science 316, 1212.1216.

Cheng, Z.N., Shu, Y., Liu, Z.Q., Wang, L.S., Ou-Yang, D.S., Zhou, H.H., 2001. Role of cytochrome P450 in estradiol metabolism in vitro. Acta Pharmacologica Sinica 22, 148–154.

Cribb, A.E., Knight, M.J., Dryer, D., Guernsey, J., Hender, K., Tesch, M., Saleh, T.M., 2006. Role of polymorphic human cytochrome P450 enzymes in estrone oxidation. Cancer Epidemiology and Prevention Biomarkers 15, 551–558.

Gaser, C., Volz, H.-P., Keibel, S., Riehemann, S., Sauer, H., 1999. Detecting structural changes in whole brain based on nonlinear deformations–Application to schizophrenia research. NeuroImage 10, 107–113.

Goldstein, J.M., Seidman, L.J., Horton, N.J., Makris, N., Kennedy, D.N., Caviness, V.S., Faraone, S.V., Tsuang, M.T., 2001. Normal sexual dimorphism of the adult human brain assessed by in vivo magnetic resonance imaging. Cereb. Cortex 11, 490–497.

Gur, R.C., Turetsky, B.I., Matsui, M., Yan, M., Bilker W Hughett, P., Gur, R.E., 1999. Sex differences in brain gray and white matter in healthy young adults: Correlations with cognitive performance. J. Neurosci. 19, 4065–4072.

Holmes, A.P., Blair, R.C., Watson, J.D.G., Ford, I., 1996. Nonparametric analysis of statistic images from functional mapping experiments. J. Cereb. Blood Flow Metab. 16, 7–22.

Ingelman-Sundberg, M., Persson, A., Jukic, M.M., 2014. Polymorphic expression of CYP2C19 and CYP2D6 in the developing and adult human brain causing variability in cognition, risk for depression and suicide: The search for the endogenous substrates. Pharmacogenomics 15, 1841–1844.

Jiang, R., Yamaori, S., Takeda, S., Yamamoto, I., Watanabe, K., 2011. Identification of cytochrome P450 enzymes responsible for metabolism of cannabidiol by human liver microsomes. Life Sci. 89, 165–170.

Jukic, M., Opel, N., Ström, J., Carrillo-Roa, T., Miksys, S., Novalen, M., Renblom, A., Sim, S.C., Peñas-Lledó, E.M., Courtet, P., Llerena, A., Baune, B.T., de Quervain, D.J., Papassotiropoulos, A., Tyndale, R.F., Binder, E.B., Dannlowski, U., Ingelman-Sundberg M,, 2016. Elevated CYP2C19 expression is associated with depressive symptoms and hippocampal homeostasis impairment. Mol. Psychiatry 22, 1155–1163.

Justenhoven, C., Hamann, U., Pierl, C.B., Baisch, C., Harth, V., Rabstein, S., Spickenheuer, A., Pesch, B., Brünung, T., Winter, S., Ko, Y.D., Brauch, H., 2009. CYP2C19*17 is associated with decreased breast cancer risk. Epidemiology 115, 391–396.

Laakso, A., Vilkman, H., Bergman, J., Haaparanta, M., solin, O., Syvälahti, E., Salokangas, R.K.R., Hietala, J., 2002. Sex differences in striatal presynaptic dopamine synthesis capacity in healthy subjects. Biol. Psychiatry 52, 759–763.

Lenroot, R.K., Giedd, J.N., 2010. Sex differences in the adolescent brain. Brain Cognition 72, 46–55.

McCarthy, M.M., 2008. Estradiol and the developing brain. Physiol. Rev. 88, 91–134.

Munro, C.Y., McCaul, M.E., Wong, D.F., Oswald, L.M., Zhou, Y., Brasic, J., Kuwubara, H., Kumar, A., Alexander, M., Ye, W., Wand, G.S., 2009. Sex differences in striatal dompamine release in healthy adults. Biol. Psychiatry 59, 966–974.

Mwinyi, J., Cavaco, I., Pedersen, R.S., Persson, A., Burkhardt, S., Mkrtchian, S., Ingelman-Sundberg, M., 2010. Regulation of CYP2C19 expression by estrogen receptor a: Implications for estrogen-dependent inhibition of drug metabolism. Mol. Pharm. 78, 886–894.

Patenaude, B., Smith, S.M., Kennedy, D., Jenkinson, M., 2011. A Bayesian model of shape and appearance for subcortical brain. NeuroImage 56, 907–922.

Pedersen, R.S., Noehr-Jensen, L., Brosen, K., 2013. Inhibitory effects of oral contraceptives on CYP2C19 activity is not significant in carriers of the CYP2C19*17 allele. Clin. Exper. Pharmacol. Physiol. 40, 683–688.

Persson, A., Ingelman-Sundberg, M., 2014. Pharmacogenomics of cytochrome P450 dependent metabolism of endgenous compounds: Implications for behavior, psychopathology and treatment. J. Pharmacogenom. Pharmacoproteomics 5, 2.

Persson, A., Sim, S.C., Virding, S., Onishchenko, N., Schulte, G., Ingelman-Sundberg, M., 2013. Decreased hippocampal volume and increased anxiety in a transgenic mouse model expressing the human CYP2C19 gene. Mol. Psychiatry 19, 733–714.

Pohjalainen, T., Rinne, J.O., Nagren, K., Syvälahti, E., Hietala, J., 1998. Sex differences I the striatal dopamie D2 receptor binding characteristics in vivo. Am. J. Psychiatry 155, 768–773.

Rijpkema, M., Everaerd, D., van der Pol, C., Franke, B., Tendolkar, I., Fernández, G., 2012. Normal sexual dimorphism in the human basal ganglia. Hum. Br. Mapping 33, 1246–1252.

Schroth, W., Antoniadou, L., Fritz, P., Schwab, M., Muerdter, T., Zanger, U.M., Simon, W., Eichelbaum, M., Brauch, H., 2007. Breast cancer treatment outcome with adjuvant tamoxifen relative to patient CYP2D6 and CYP2C19 genotypes. Journal of Clinical Oncology 33, 5187–5193.

Sheline, Y.I., 2011. Depression and the hippocampus: Cause or effect? Biol. Psychiatry 70, 308–309.

Sim, S.C., Risinger, C., Dahl, M.L., Eklillu, E., Christensen, M., Bertilssonn, L., Ingelman-sundberg, M., 2006. A common novel CYP2C19 gene variant causes ultrarapid drug metabolism relevant for the drug response to proton pump nhibitors and antidepressants. Clin. Pharmacol. Ther. 79, 103–113.

Stingl, J.C., Brockmöller, J., Viviani, R., 2013. Genetic variability of drug-metabolizing enzymes: The dual impact on psychiatric therapy and regulation of brain function. Mol. Psychiatry 18, 273–287.

Stingl, J.C., Jukic, M., Tyndale, R., Steffens, M., Paul, A.M., Viviani, R., 2016. Effects of CYP2C19 polymorphism on voxel-based morphometry. Human Brain Mapping Annual Meeting, Geneva, June 29 2016.

Viviani, R., Beschoner, P., Ehrhard, K., Schmitz, B., Thöne, J., 2007. Non-normality and transformations of random fields, with an application to voxel-based morphometry. NeuroImage 35, 121–130.

Viviani, R., Stöcker, T., Stingl, J.C., 2017. Multimodal FLAIR/MPRAGE segmentation of cerebral cortex and cortical myelin. NeuroImage 152, 130–141.

